# Pix2Path: Integrating Spatial Transcriptomics and Digital Pathology with Deep Learning to Score Pathological Risk and Link Gene Expression to Disease Mechanisms

**DOI:** 10.1101/2024.08.18.608468

**Authors:** Xiaonan Fu, Yan Chen

## Abstract

Spatial transcriptomics (ST) provides high-resolution mapping of gene expression within tissues, and integrating ST with digital pathology can offer unprecedented insights into the molecular mechanisms underlying various diseases. However, existing methods primarily focus on aligning these two distinct datasets, often neglecting the causal connections between spatial gene activity and pathological phenotype. We introduce Pix2Path, a deep learning-based approach utilizing conditional generative adversarial networks (cGANs), to bridge the gap between spatial transcriptomics and digital pathology. Pix2Path can process data from various spatial transcriptomics (ST) technologies, assess pathological risk scores across different conditions, and supports a leave-one-out spatial in silico gene perturbation strategy. As demonstrated in AD Aβ plaques pathology, this approach allows to link gene expression changes to tissue morphology and pathology without relying on predefined conditions, providing a new perspective on understanding disease mechanisms.

## Main

Digital pathology primarily focuses on the morphological aspects of tissues, often overlooking the underlying molecular mechanisms driving disease processes. In contrast, the advancement of spatial transcriptomics (ST) (*1, 2*), provide subcellular-resolution precise mapping of whole transcriptomics within the native tissue context (*3, 4*). These methods offer insights into the spatial organization of gene activity and its relationship to disease pathology.

The application of deep learning models in digital pathology is rapidly revolutionizing the field (*5, 6*), enabling automated tissue analysis, disease detection, and classification at an unprecedented scale and accuracy (*7*). Deep learning models excel in tasks such as tissue grading (*8*), virtual stain transformation (*9*) and tumor heterogeneity quantification (*7*), making them invaluable in modern diagnostic workflows. With the integration of spatial omics data and deep learning tools, there is an opportunity to assess pathological risk more effectively, potentially facilitating earlier diagnosis.

Previous computational approaches have mainly focused on spatially aligning morphology images with spatial datasets (*10*), or aligning datasets across different spatial transcriptomics technologies (*11-15*). Deep learning model has been successfully to integrate spatial transcriptomics to infer the survival outcome from whole slide digital image (*7*). However, A significant challenge remains in directly inferring gene regulatory networks from spatial gene expression data and linking these networks to observed pathological changes and access the pathological risk.

To address these challenges, we present an approach called Pix2Path (**Fig. 1a**) that builds on recent developments in a deep learning-based model conditional generative adversarial network (cGAN) (*16*) to bridge the gap between spatial transcriptomics and digital pathology. Pix2Path is amenable to data from various ST technologies for which a corresponding pathology image is available. The trained model enables the estimation of pathological scores across different conditions. Pix2Path is further able to perform spatial in silico gene perturbations. Unlike distance-based methods for identifying pathology-associated genes (*17*), it facilitates an unbiased exploration of how gene expression variations impact tissue morphology and pathology. By linking spatial transcriptomics data to digital pathology, Pix2Path provides a novel approach to studying the molecular underpinnings of diseases, particularly those with complex spatial and morphological characteristics.

**Fig. 1.**
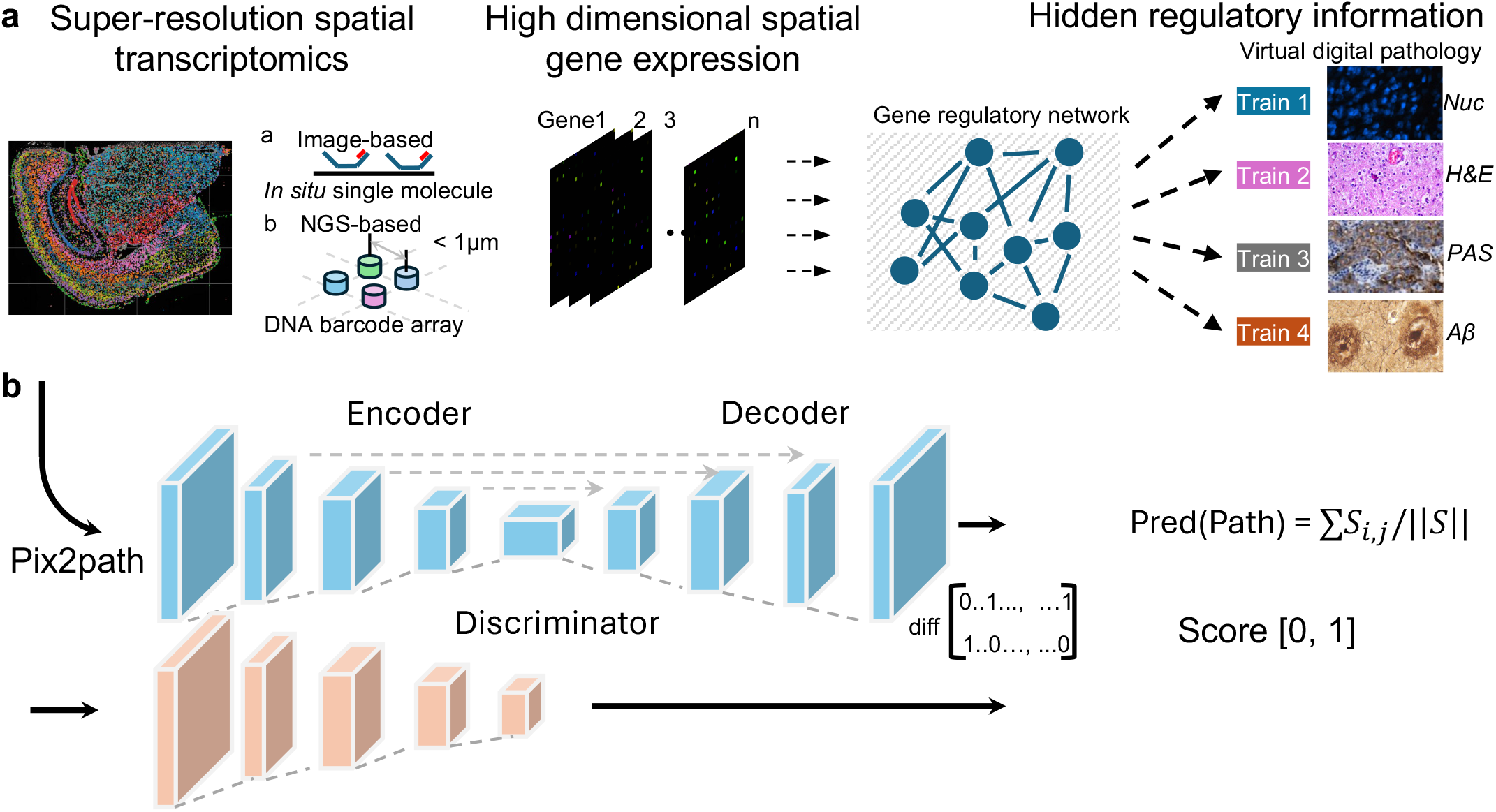
Pix2Path inferring virtual digital pathology image from spatial transcriptomic data. **a**, Utilizing high-resolution spatial transcriptomics data, which includes imaging-based single-molecule sequencing and NGS-based DNA barcoding methods, we pair these with digital pathology images as inputs. A deep learning model is then trained to uncover the hidden gene regulatory networks, enabling it to generate virtual pathology images that accurately replicate the observed tissue architecture. **b**, The Pix2Path model is an adaptation of the Pix2Pix architecture, featuring a generator based on a U-Net design and a discriminator modeled as a convolutional PatchGAN classifier. The synthesized images produced by the model are normalized and interpreted as pathology scores.

## Results

### Pix2Path inferring virtual digital pathology image from spatial transcriptomic data

Pix2Path leverages high-resolution (<1*µ*m) spatial transcriptomics data (6, 7) alongside paired digital pathology images to uncover hidden networks (**Fig. 1a**). The model learns these connections in two key steps (**Fig. 1a**). First, a U-Net model extracts both local and global features from the spatial transcriptomics data to infer a potential pathological outcome (**Fig. 1b**). Next, a discriminator network with fully connected layers compares the inferred outcome to the actual pathological observations, evaluating it through a defined loss function.

To test the network, the model was first trained on a segmentation-relevant task. Cell segmentation poses a significant challenge in spatial omics for single-cell level analysis, as it typically relies on pre-scanned nucleus images (18, 19). The Pix2Path model successfully predicted nucleus masks using the defined gene sets (**Fig. S1a, Table S2**), generate virtual hematoxylin and eosin (H&E) images (**Fig. S1b**). We further applied Pix2Path to various high-resolution spatial transcriptomics platforms, including a mouse Alzheimer”s disease brain dataset profiled with Stereo-seq featuring nucleus and H&E staining (*20*), a mouse brain dataset from the 10x Genomics Xenium platform with nucleus and Aβ plaque staining, as well as datasets from NanoString CosMx and Pixel-seq. To assess performance, the model was trained separately for each platform. Our results demonstrate that Pix2Path can accurately recreate virtual pathology images from spatial transcriptomics data (**Fig. S1c, d**).

### Pix2Path provide accurate Aβ plagues risk prediction

Aβ plaques are one of the pathological hallmarks of AD; additionally, amyloid itself exerts toxic effects on cells and promotes neuroinflammation (*21*). To assess the performance and robustness of Pix2Path, we applied it to a Xenium AD dataset that includes three time points: 2.5, 5.7, and 13.2 months of age for WT mice, and 2.5, 5.7, and 17.9 months of age for TgCRND8 mice. These time points correspond to different stages of disease progression, specifically mild, moderate, and advanced Aβ deposition (**Fig. 2a**). On average, 58,619 cells are detected per section. The median transcripts per cell were around 205 (**Fig. 2b**). We selected data from 17.9 months of TgCRND8 for the training, the Aβ plagues image were first determine the background threshold (**Fig. S2a**), clean, and normalized to range of [0,1]. The predicted Aβ plagues image clearly shows agreement with the ground truth (**Fig. 2c**).

**Fig. 2.**
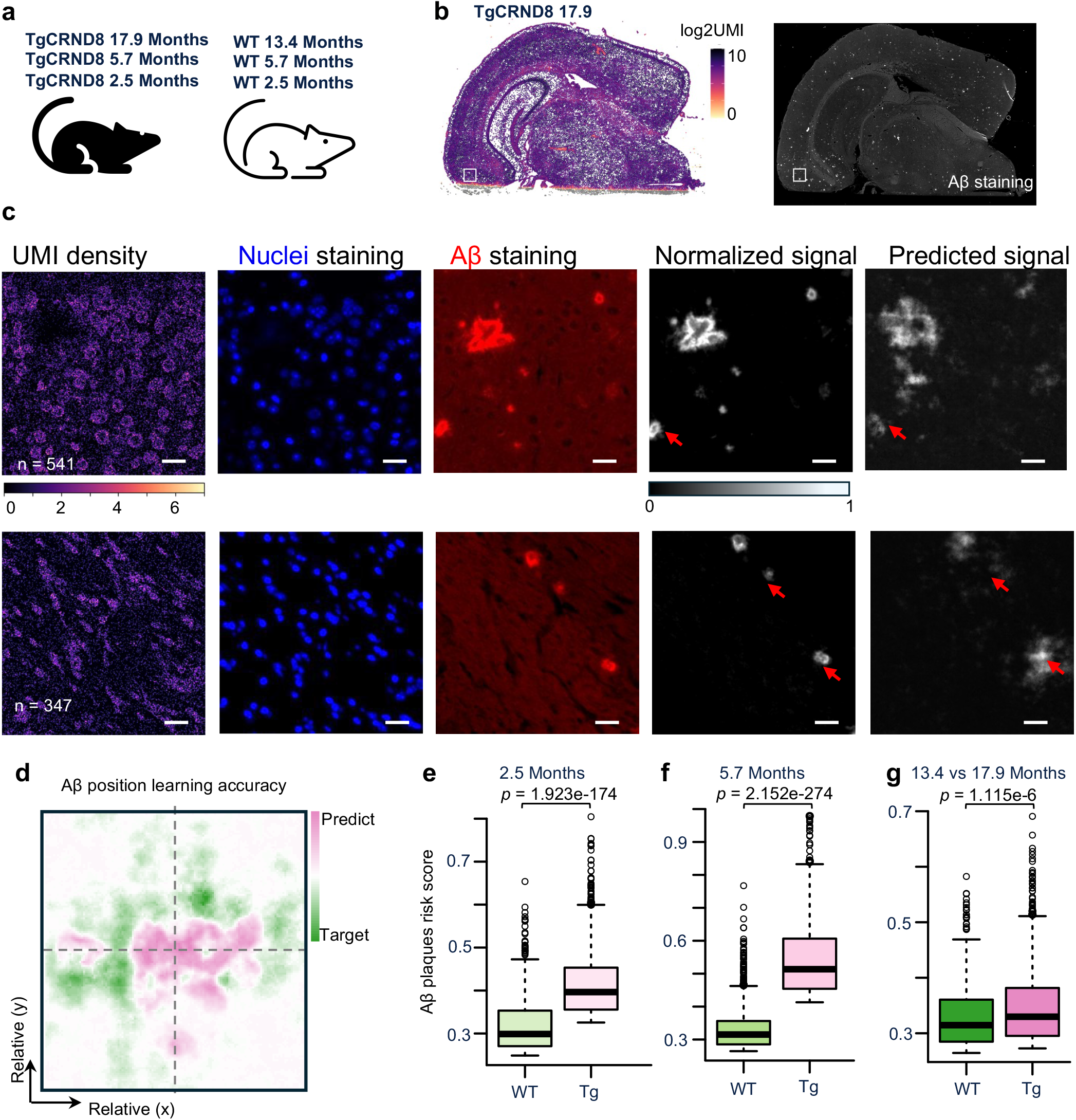
Pix2Path provide acurate Aβ plagues risk prediction. **a**, The dataset used to train the Pix2Path model in AD. The Xenium mouse AD datasets were generated from wild-type (WT) and a well-characterized transgenic AD model (TgCRND8) mice. **b**, The 10x Genomics Xenium platform generate single molecule data alongside immunofluorescence (IF)-labeled Aβ protein plaques. **c**, Images shows Pix2Path predicted results on Aβ plaques pathology. **d**, Heatmap shows the positional accuracy of predicted Aβ plaques. **e, f and g**, Boxplot shows the total Aβ plaques risk score calculated from each sample.

In this evaluation, we also compared the Aβ plaque spots where the ground truth and predicted regions were expected to align (see Methods for details). Specifically, for each Aβ plaque spot in the ground truth image, we identified the geometric center and extracted a 200 *µ*m x 200 *µ*m square area for comparison. As shown, the predicted regions closely matched the centers of the Aβ plaques in the ground truth (**Fig. 2d**). Additionally, we define the Aβ plagues risk score (see Methods for details) to summarize the overall effect from each prediction. the overall pathological score per image exhibited a strong correlation with the ground truth (**Fig. S2b**, R > 0.9). We then applied the model to data from other time points to calculate the Aβ plaque risk score. As expected, samples from the transgenic AD mouse model show a significantly higher Aβ plaque risk score at 2.5 and 5.7 months (**Fig. 2e, f**). Interestingly, the predicted Aβ plaque risk score at 13.4 months for WT mice compared to 17.9 months for AD mice shows a decreasing trend, which suggests an raised risk as the mice age (**Fig. 2g**).

In summary, these results demonstrate that Pix2Path is capable of generating reliable virtual pathological images and can be effectively used to predict pathological risk scores, making it a valuable tool for early diagnosis.

### In silico spatial gene perturbation by Pix2Path predict Aβ plaque genes

Spatial transcriptomics is a powerful tool for studying cell-type expression and interactions, and high-resolution methods enable the characterization of molecular heterogeneity within individual cells. However, pathological differential expression analysis using spatial transcriptomics often depends on spatial coordinates and manually defined thresholds to differentiate between proximal and distal regions for determining disease-associated gene lists (*22, 23*). This approach may miss relevant factors that are located at longer distances.

Although Pix2Path is designed to predict virtual pathological images, its high-resolution spatial gene expression data and underlying network structure also provide the opportunity to manipulate the input gene expression matrix. This enables an unbiased assessment of each gene”s potential contribution to the pathological phenotype (**Fig. 3a**). To obtain a robust and statistically significant gene list, we introduced randomized genes as negative controls (see Methods for details). In this test, we calculated the summarized Aβ plaque offset score across the entire dataset. The Aβ plaque offset score per gene exhibited an interesting variation pattern. While the negative controls were within a range of [-0.25, 0.25], the detected Aβ plaque relevant genes, agree with previously identified as amyloid plaque-induced genes (PIGs)(*17*) (**Fig. S3a, Fig. 3b**), showed distinct patterns. For example, the upper offset genes Hexb and Cst3 are markers of glial cell states, while Apoe is involved in Aβ metabolism. Conversely, the lower offset genes Ctsd and B2m are crucial for antigen processing and presentation, and Gfap is important for the cytoskeletal structure in glial cells. Notably, the upper offset genes appear to be relevant at a distance, whereas the lower offset genes are proximity-relevant (**Fig. 3d**). Upon this observation, we tested the effect of different kernel size on the model (**Fig. S3b**).

**Fig. 3.**
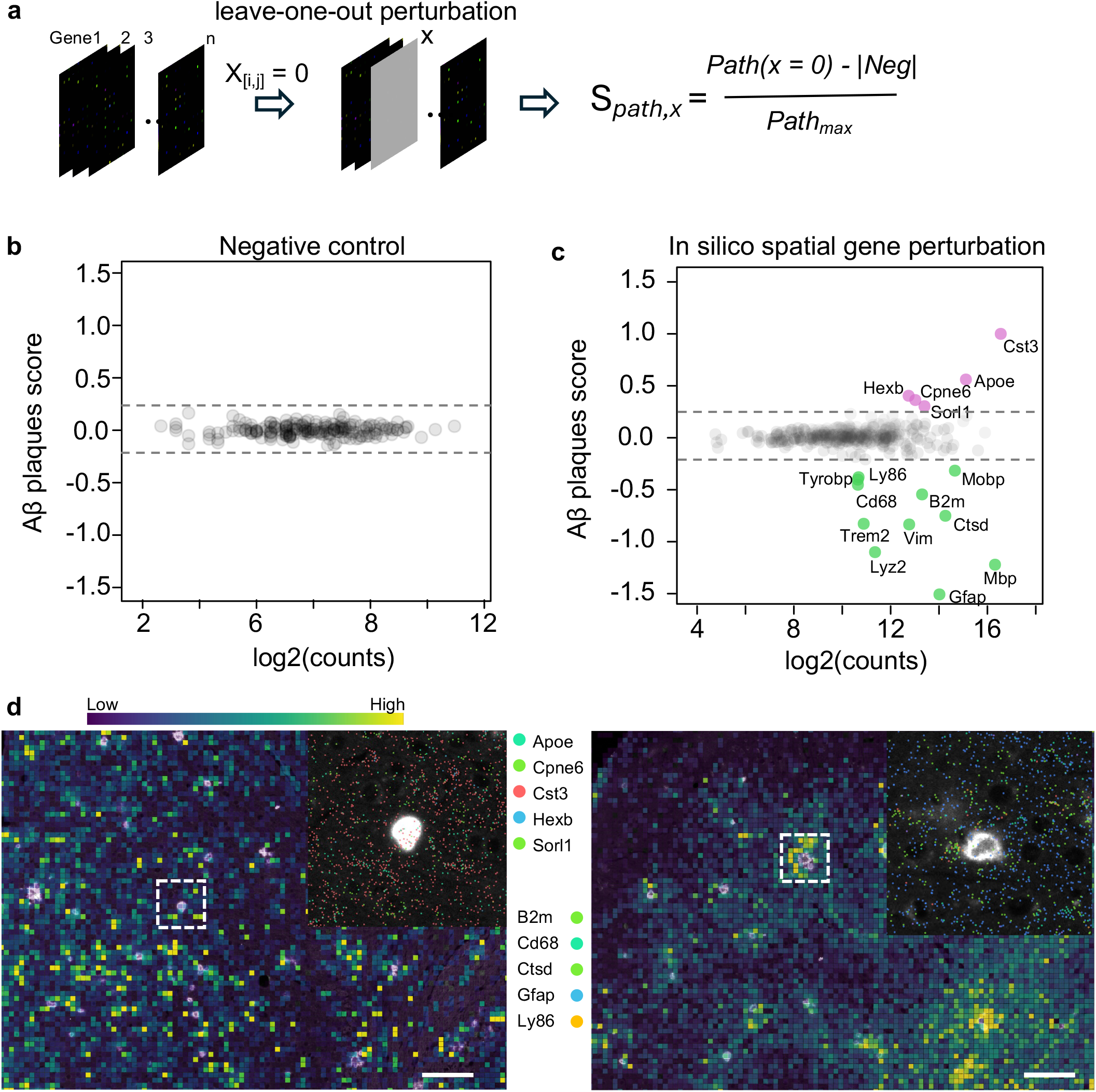
Pix2Path enable spatial in silico gene perturbation. **a**, The strategy for performing spatial in silico gene perturbation involves a “leave-one-out” approach to assess the impact on the resulting pathology outcome. **b**, The scatter plot displays the distribution of Aβ plaque score offsets (y-axis) against the relative expression of each gene (x-axis). The left panel describe 194 negative random controls, the right panel includes all the 347 genes in the assay (**c**). Offset genes are labelled in color. **d**. Spatial visualization of the upper offset gene sets (left) and lower offset gene sets (right).

These results indicate that Pix2Path can greatly improve the unbiasedly identification of pathology-related genes using high-resolution spatial transcriptomics data.

## Discussion

Recent advancements in spatial transcriptomics offer sufficient resolution and depth to develop deep learning models that can uncover hidden regulatory networks, potentially linking omic-scale gene expression to pathological phenotypes. High-resolution, high-dimensional spatial transcriptomics (ST) data contain rich information, making deep learning an effective strategy for extracting and interpreting these complex patterns. However, existing studies have primarily focused on the integration of multiple datasets (*10, 15*), alignment of images to omic data (*11-13*) and image-assisted cell segmentation (*18, 19*). These approaches do not fully exploit the interconnected gene expression data to explore its pathological relevance.

Pix2Path harnesses high-resolution spatial transcriptomics data to infer virtual digital pathology images, bridging the gap between spatial gene expression and pathological observations. By utilizing a U-Net model to extract both local and global features and a discriminator network to compare these features with actual pathological data, Pix2Path effectively generates accurate virtual pathology images. This capability was demonstrated across multiple high-resolution spatial transcriptomics platforms, including datasets from mouse Alzheimer”s disease brains, where Pix2Path successfully predicted nucleus masks and virtual hematoxylin and eosin (H&E) images. The model”s robust performance across different datasets highlights its potential to recreate detailed pathological images and provides reliable insights into disease progression.

Furthermore, Pix2Path”s application to Aβ plaque risk prediction showcases its utility in identifying pathology-related genes. By analyzing data from various stages of Alzheimer”s disease in mouse models, Pix2Path accurately predicted Aβ plaque risk scores, demonstrating strong correlation with ground truth observations. The model”s ability to assess the contribution of individual genes to the pathological phenotype—without relying on predefined spatial thresholds—enhances its capacity to identify long-distance relevant factors and provide unbiased gene assessments. This capability is crucial for advancing our understanding of disease mechanisms and improving early diagnostic strategies.

## Methods

### Spatial transcriptomics dataset

We applied Pix2Path to spatial transcriptomics (ST) datasets generated by various platforms, including 10x Xenium, Stereo-seq, NanoString CosMx, and Pixel-seq (see Supplementary Table S1 for details, GEO accession number). The training dataset comprised 6 human tissue sections, 10 sections from App^NL-G-F^ mice along with paired wildtype controls, and six sections from the TgCRND8 Alzheimer”s disease mouse model. We obtained the raw datasets, which included spatial coordinates, corresponding gene expression data, and the paired pathology images.

### Data preprocessing

In the Stereo-seq and Pixel-seq ST datasets, we first removed spots located outside the main tissue area. The raw gene expressions were then log-transformed and normalized according to library size using the SCANPY package (*24*). We selected the top 500 highly variable genes to prepare the input matrix, each layer represents a single gene. The paired pathology image was aligned and normalized to a [0, 1] range relative to the background signal. The prepared matrix was then cropped into appropriately sized patches (512 x 512), and patches with a pathology score below 0.1 were excluded. For the 10x Xenium and NanoString CosMx datasets, all genes were included in the input (see Supplementary Note 1).

### Training of Pix2Path transformation network

An overview of the model is illustrated in Fig. 1b. The model consists of two components: a generator based on a U-Net architecture (encoder and decoder) and a discriminator implemented as a convolutional PatchGAN classifier. Both the generator and discriminator utilize modules structured in the form of convolution, followed by BatchNorm, and then ReLU. Different from previous spatial transcriptomics preprocessing methods (*18*), the encoder maps each gene layer to a hidden representation layer without binned from neighborhood expression. The paired pathology image was transformed to a dimension that matches pixel-to-pixel with the input data.

The loss function of a Pix2Path can be expressed as:

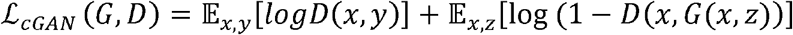

where generator G tries to minimize this objective function against discriminator D. Gaussian noise z were generated as an input to the generator in addition to x.

### Spatial in silico gene perturbation using Pix2Path model

The single-layer input per gene in the Pix2Path model allows for testing the individual contribution of each gene to the final output. Specifically, during pathological outcome training, the layer corresponding to each gene was set to zero at every position while the other gene layers remained active, generating a predicted image. A control layer with random assignments was used as a comparison. The pathology offset score was then calculated by:

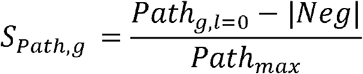

Finally, the pathology offset score was summarized and normalized for further analysis.

### Implementation details

The image alignment was performed in squidy version 1.5.0 package(*25*). The neural networks were trained and implemented using Python version 3.10.1 with TensorFlow version 1.8.0. The timing was measured on a Linux workstation computer with two Nvidia GeForce GTX 1080 Ti GPUs, 128GB of RAM, and an Intel I9-7900X CPU.

### Statistical analysis

A Kolmogorov–Smirnov test was used to compare the pathology score between two samples in R. For all tests, a P value of 0.05 or less was significant.

## Data availability

The data will be released once the paper be accepted.

## Code availability

Pix2Path code, documentation and tutorials are available at GitHub: https://github.com/frankXiaonan/Pix2Path.

